# Nonlinear Interaction Decomposition (NID): A Method for Separation of Cross-frequency Coupled Sources in Human Brain

**DOI:** 10.1101/680397

**Authors:** Mina Jamshidi Idaji, Klaus-Robert Müller, Guido Nolte, Burkhard Maess, Arno Villringer, Vadim V. Nikulin

## Abstract

Cross-frequency coupling (CFC) is a phenomenon through which spatially and spectrally distributed information can be integrated in the brain. There is, however, a lack of methods decomposing brain electrophysiological data into interacting components. Here, we propose a novel framework for detecting such interactions in Magneto- and Electroencephalography (MEG/EEG), which we refer to as Nonlinear Interaction Decomposition (NID). In contrast to all previous methods for separation of cross-frequency (CF) sources in the brain, we propose that the extraction of nonlinearly interacting oscillations can be based on the statistical properties of their linear mixtures. The main idea of NID is that nonlinearly coupled brain oscillations can be mixed in such a way that the resulting linear mixture has a non-Gaussian distribution. We evaluate this argument analytically for amplitude-modulated narrow-band oscillations which are either phase-phase or amplitude-amplitude CF coupled. We validated NID extensively with simulated EEG obtained with realistic head modeling. The method extracted nonlinearly interacting components reliably even at SNRs as small as −15 (dB). Additionally, we applied NID to the resting-state EEG of 81 subjects to characterize CF phase-phase coupling between alpha and beta oscillations. The extracted sources were located in temporal, parietal and frontal areas, demonstrating the existence of diverse local and distant nonlinear interactions in resting-state EEG data.

## 1. Introduction

Oscillatory neuronal activity has been associated with almost all brain operations including sensory, motor and cognitive processes (Buzsáki and Draguhn, 2004). In humans, these oscillations can be measured with magneto- and electroencephalography (MEG/EEG), where the frequency content is classically divided into specific frequency bands, namely *δ* (0.5-4Hz), *θ* (4-8Hz), *α* (8-12Hz), *β* (12-25Hz), *γ* (25-70Hz)^1^. Each frequency band has been associated with specific functional roles. For example, alpha oscillations are known to be relevant for attention/sensory processing (Groppe et al., 2013; Klimesch, 2012), while beta-band activity is primarily associated with sensorimotor processing (Bayraktaroglu et al., 2011; Kilavik et al., 2013; Klimesch, 2012; Salmelin and Hari, 1994). While specific neuronal operations can be carried out by oscillations in distinct frequency bands, there should be neuronal mechanisms integrating such spatially and spectrally distributed processing (Palva et al., 2005). In this way neuronal communications can be considerably enriched via coupling of neuronal oscillations within one frequency band (Engel and Fries, 2010; Fries, 2015) as well as between different frequency bands. Various types of cross-frequency (CF) interactions among neural oscillations, namely phase-phase, amplitude-amplitude, phase-amplitude coupling have been observed in human electrophysiological recordings (e.g. MEG/EEG) (Canolty and Knight, 2010; Jensen and Colgin, 2007; Nikulin and Brismar, 2006; Palva et al., 2005) and have been linked to diverse perceptual and cognitive processes (Canolty and Knight, 2010; Fell and Axmacher, 2011; Hyafil et al., 2015; Palva et al., 2005; Sauseng et al., 2008; Siebenhühner et al., 2016). In this study, we focus on the extraction of these interactions from multi-channel MEG/EEG. While the novel approach introduced here is applicable to different types of CFC, a special emphasis is dedicated to phase-phase coupling for the following reasons.

The phase of neuronal oscillations is known to represent the timing of the firing of a neuronal population generating the oscillation (Fries, 2009, 2015; Palva et al., 2005; Siegel et al., 2012), while its amplitude reflects the strength of local spatial synchronization (Siegel et al., 2012). The interaction of the activities of distinct neuronal populations is manifested in the locking of phase/amplitude of the observed oscillations. Phase-phase coupling is a type of CFC that operates with millisecond precision for both oscillations (Fell and Axmacher, 2011; Palva et al., 2005; Siegel et al., 2012) and investigating it with MEG/EEG recordings can provide a unique possibility to study synchronization of the spiking of distinct neuronal populations non-invasively (Palva and Palva, 2018).

A number of previous studies have investigated CF phase synchronization in sensor-space (Darvas et al., 2009; Nikulin and Brismar, 2006; Palva et al., 2005; Tass et al., 1998). However, volume conduction does not allow the disentanglement of individual components. In order to resolve this issue, some previous studies have investigated the phase synchrony in the source-space using inverse modelling (Siebenhühner et al., 2016; Tass et al., 2003). Yet, source-space analysis is computationally exhausting and source reconstruction methods are ill-posed, which may lead to inconsistent outcomes (Mahjoory et al., 2017). On the other hand, due to a linear mapping of the neuronal source signals to the sensors, multivariate methods can increase the signal-to-noise ratio (SNR) and accuracy of localizing the neuronal activity (Parra et al., 2005). At the same time, these methods alleviate the problem of multiple testing in sensor- or source-space analysis. While most of the multivariate source-separation methods focus on the extraction of independent sources (e.g. independent component analysis - ICA), there are only a few studies utilizing multivariate methods to extract dependent sources from the electrophysiological recordings of the human brain (Chella et al.; Cohen, 2017; Dähne et al., 2014; Nikulin et al., 2012; Volk et al., 2018). These methods optimize a contrast function of the desired type of coupling. However, we show that the coupling can be reflected in the statistical properties of the signal constructed through the linear mixing of nonlinearly coupled processes. We refer to our method as Nonlinear Interaction Decomposition (NID).

The rest of the manuscript is organized as follows. In section 2 we discuss the distribution of a linear mixture of cross-frequency coupled oscillations, which represents the main idea of this work. Section 3 is dedicated to explaining the proposed method (NID) and its algorithmic steps. In section 4 the experimental data and the analysis/testing approaches are described. The results of applying NID to simulated as well as resting-state EEG data are presented in section 5. Finally, a discussion and conclusion are provided in the last section.

## 2. Linear mixture of non-linearly coupled oscillations

The oscillations observed in the brain are amplitude-modulated narrow-band signals. The envelope of these oscillations fluctuate much slower than the carrier frequency and shows long-range temporal correlation (Linkenkaer-Hansen et al., 2001). This property leads to the phenomenon that the distribution of the signal does not deviate strongly from Gaussian distribution (Hyvärinen et al., 2010). Therefore, each oscillation (with T time-samples) can be assumed as T realization of a Gaussian random variable.

Any linear combination of independent or linearly dependent Gaussian random variables is Gaussian. However, the distribution of a linear mixture of nonlinearly dependent Gaussian random variables is not necessarily Gaussian. While linear dependency of two oscillations means that there is a within-frequency coupling between them, nonlinear dependency relates to CFC. The core idea of the current work comes from our observation that it is possible to find weights for two CF coupled oscillations such that the resulting mixed signal has a distribution being more non-Gaussian than the distribution corresponding to each individual oscillations. We have analytically proved this observation for phase-phase and amplitude-amplitude coupled amplitude-modulated narrow-band signals (supplementary text, section 1). Figure 1 illustrates the working principle of NID. Note that we assess the non-Gaussianity of a random variable by means of kurtosis, skewness, or 5th order moments, all of which are zero for Gaussian random variables.

**Figure 1:**
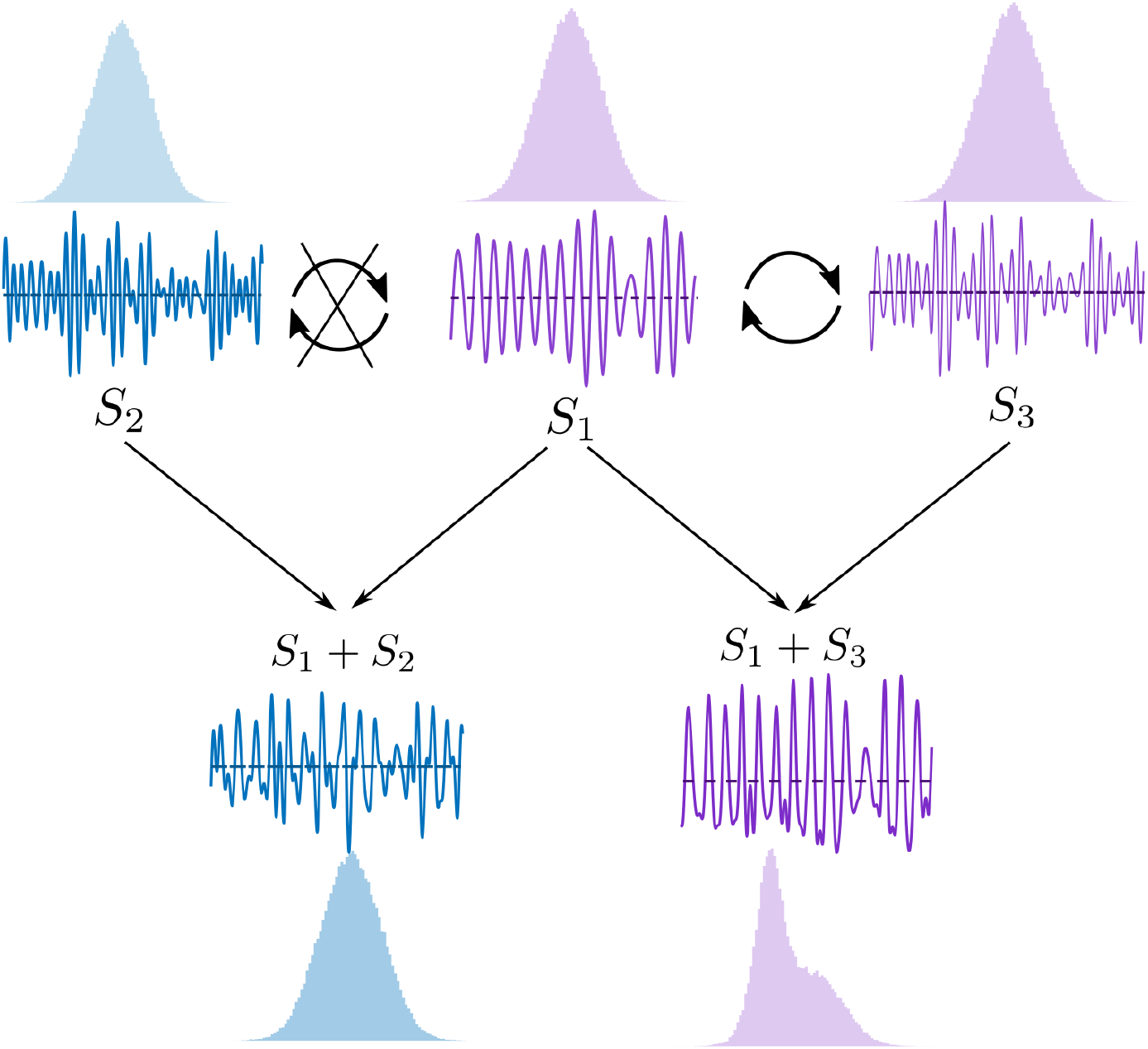
The core idea of the proposed method (NID). The signals in the bottom of the figure are linear mixtures signals of *S*_1_ and *S*_*i*_, *i* = 2, 3. The shaded histograms are the amplitude distributions of the signals. *S*_1_ and *S*_3_ are CF phase-phase coupled, while *S*_1_ and *S*_2_ are independent signals with different central frequencies. The distribution of the linear mixture is non-Gaussian only in the case of the existence of nonlinear dependency between the two amplitude-modulated narrow-band oscillations.

## 3. Method

### 3.1. Notation

We use boldface lower-case letters (e.g. **x**) to denote vectors, while bold-face capital letters (e.g. **X**) are used for matrices. Regular letters, (e.g. *x*), indicate scalars. Vectors are used to denote the time series of a signal or spatial filters/activation patterns. Matrices are used to denote the concatenation of vectors. The operators [., .] and [.; .] stand for horizontal and vertical concatenation of two matrices respectively.

### 3.2. Measuring Cross-Frequency Coupling

Depending on the type of the coupling, there are different measures to quantify CFC. In this paper, we worked with phase-phase and amplitude-amplitude coupled oscillations. As described below, the phase locking value (PLV) was used for measuring phase-phase coupling, while amplitude-amplitude coupling was quantified with the envelope correlation. Both of these measures are calculated from the instantaneous phase and amplitude of oscillations, which are computed as the phase and magnitude of the complex analytic signal based on the Hilbert transform.

#### Phase-phase coupling

Oscillations with frequencies *f*_*n*_ and 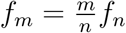, *n*, *m* ∈ ℕ are called phase-locked |*m*Φ_*n*_(*t*) − *n*Φ_*m*_(*t*)| < *const*, where Φ_*n*_(*t*) and Φ_*m*_(*t*) define the instantaneous phases of the two oscillations at *f*_*n*_ and *f*_*m*_ respectively. To quantify *n* : *m* phase-phase coupling, phase-locking value (PLV) is widely used (Palva et al., 2005; Sauseng et al., 2008; Scheffer-Teixeira and Tort, 2016; Siebenhühner et al., 2016) and it is defined as 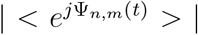, where Ψ_*n,m*_(*t*) = (*m*Φ_*n*_(*t*) − *n*Φ_*m*_(*t*)), < . > stands for computation of the mean over time samples, *j* is the imaginary number, and |.| is the absolute value operator.

#### Amplitude-amplitude coupling

In the case of amplitude-amplitude coupling, the instantaneous amplitudes of oscillations are correlated. Therefore, the correlation coefficient of the oscillations’ envelopes indicates the strength of amplitude-amplitude coupling.

### 3.3. Detection of cross-frequency coupling: problem formulation

We assume that there are *N* non-linearly coupled pairs of source signals 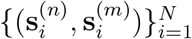 at frequencies *f*_*n*_ and *f*_*m*_, where *f*_*n*_ = *nf*_*b*_ and *f*_*m*_ = *mf*_*b*_. *f*_*b*_ is a base-frequency relating *f*_*n*_ and *f*_*m*_ to each other. In the rest of the paper, all the criteria and equations mentioned for frequency *f*_*n*_ holds for frequency *f*_*m*_ as well. 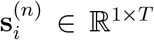 is a narrow-band source signal at *f*_*n*_, where *T* is the number of time samples. The electrical (or magnetic) activity measured at the sensors can be modeled as a linear mixture of the sources as in the following (Baillet et al., 2001; Haufe et al., 2014):

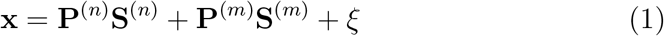

where **X** ∈ ℝ^*C*×*T*^ is the matrix of multi-channel measured signal with *C* as the number of channels. 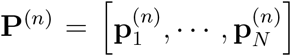. We call 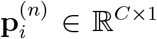 the mixing pattern of source 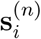. Additionally, 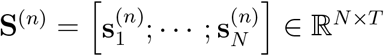 is the matrix of source signals at *f*_*n*_, which are CF coupled to sources in matrix 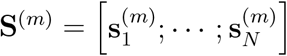. In the equation 1, *ξ* denotes the noise signal, which cannot be explained by the linear model. Note that the superscript of the sources is an indication of their frequency, e.g the superscript (*n*) in 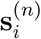 is related to the subscript *n* of *f*_*n*_. As mentioned in section 3.2, the coupling is called *n* : *m* coupling if 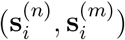 are phase-phase coupled. However, we use this notation for amplitude-amplitude coupling as well so that we can denote the frequency ratios easier.

We provide an example here. Assume that we have 2 coupled source signals in *α* and *β* frequency band, i.e. *N* = 2, *n* = 1, *m* = 2, and *f*_*b*_ = 10*Hz, f*_1_ = 10*Hz, f*_2_ = 20*Hz*. Then 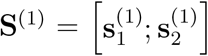 and 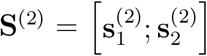 are the CF coupled source signal matrices at frequencies *f*_1_ and *f*_2_ respectively.

The main problem here is the retrieval of the sets of interacting sources and their corresponding mixing patterns from the multi-channel signal **X**. In other words, there is an additional question compared to typical source separation problems, i.e. detection of the coupling between the sources.

The above mentioned problem can be generalized to more coupled sources:for instance, triplets of coupled source signals 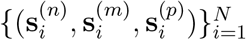. As an example, one can think of coupling of source signals with central frequencies of 10Hz (*α*), 20Hz (*β*), and 30Hz (low *γ*) source signals, which we call 1:2:3 coupling. In this manuscript, we mainly focus on the source model in equation 1 for pairs of coupled source signals. However, we test the performance of the method with simulated data for extraction of triplets of source signals with 1:2:3 coupling.

### 3.4. Nonlinear Interaction Decomposition (NID)

As discussed in section 2, the working principle of NID is that phase-phase and amplitude-amplitude coupled amplitude-modulated narrow band signals can be linearly mixed in the way that the linear mixture has non-Gaussian distribution. On the other hand, the linear mixture of independent oscillations is approximately normally distributed.

The NID algorithm consists of two main steps: first, applying a method to extract approximate estimates of the sources at *f*_*n*_ and *f*_*m*_ and second, maximizing the non-Gaussianity of the weighted sum of projected signals (refer to section 2). A block diagram of NID is depicted in figure 2.

**Figure 2:**
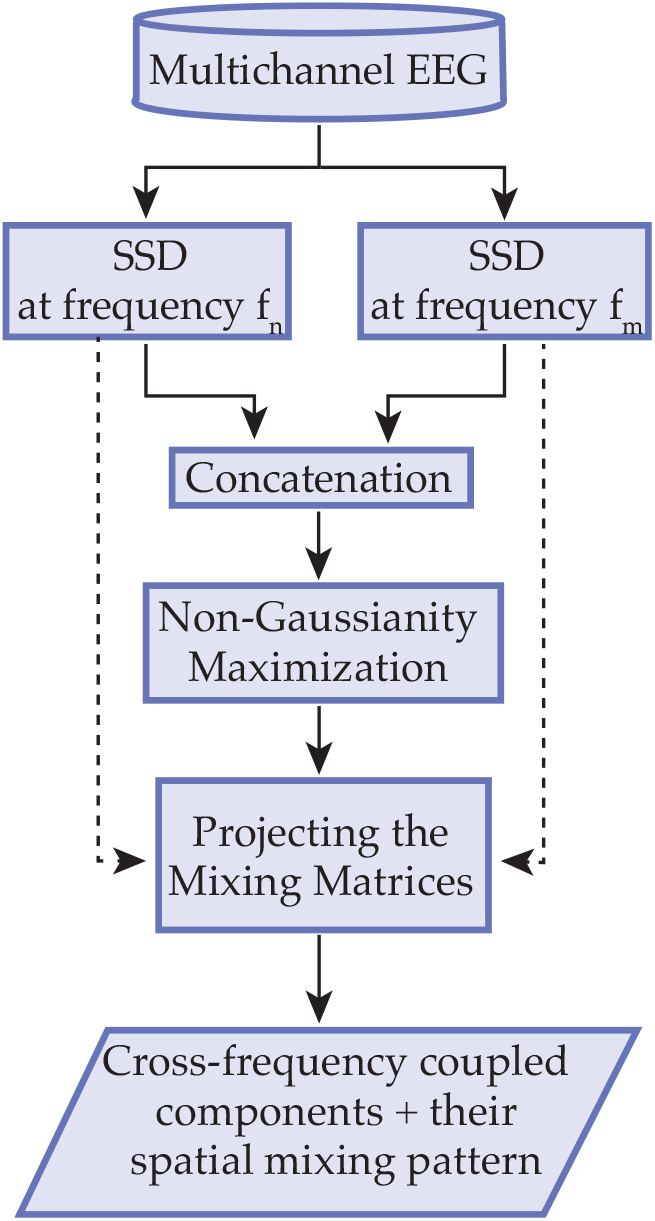
Block-diagram of NID. The input is the multi-channel mixed signal. Then SSD is applied to extract estimates of the source signals at *f*_*n*_ and *f*_*m*_. The SSD components are concatenated vertically to build an augmented matrix with its rows being estimates of source signals in the two frequency bands. The augmented matrix is then passed to a non-Gaussianity maximization (NGM) algorithm for the separation of the coupled sources. In the last step the weights computed by NGM algorithm are combined with the mixing matrices of SSD in order to build the mixing patterns of the narrow-band oscillations. In the output, the time course and the mixing patterns of the coupled sources are extracted.

In the first step, Spatio-Spectral Decomposition (SSD) (Nikulin et al., 2011) is applied to extract *N* oscillations from the multi-channel mixed signal. SSD is a method based on generalized eigenvalue decomposition, which calculates the spatial filters that maximize the SNR at the frequency band of interest. We denote the source model of SSD as 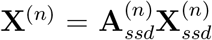, where **X**^(*n*)^ ∈ ℝ^*C*×*T*^ contains mixed, narrow-band, multi-channel signal with central frequency *f*_*n*_. 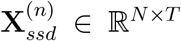 is the matrix of SSD components, and 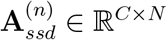 is the SSD mixing matrix.

In the second step of NID, the objective is to find a subspace, in which linear mixtures of SSD components are maximally non-Gaussian, which results in separating the cross-frequency coupled oscillations (Appendix A). This is done through applying a non-Gaussianity maximization decomposition (NGMD) on an augmented matrix 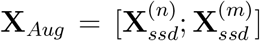. In this way, NGMD finds the linear mapping, which maximizes non-Gaussianity of the linear mixtures of the source signal estimates. The NGMD can be formulated as **X**_*Aug*_ = **A**_*NG*_**R**_*NG*_, where **R**_*NG*_ includes the linear mixtures of SSD components. 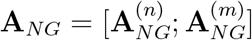 is the NGMD linear mapping and 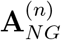 is the NGMD linear mapping corresponding to SSD components of frequency *f*_*n*_. The final mixing matrix, revealing an estimation of the activation patterns of the coupled oscillations can be computed by back-projecting 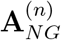 from SSD space to sensor space:

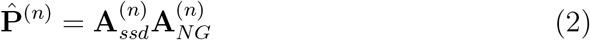

We refer the reader to Appendix A and B for more discussion on NGMD and some practical details about computing the final mixing patterns. Additionally, the MATLAB^©^ codes of the NID algorithm are publicly accessible on GitHub (https://github.com/minajamshidi/NID).

### 3.5. Statistical testing of coupling

Statistical testing has been applied in order to control for the effects of overfitting when extracting coupled components. For this purpose, the SSD components of the lower frequency were cut into one-second segments, which were then randomly permuted. The NGMD was applied to the augmented matrix of permuted SSD components of the lower frequency and SSD components of the higher frequency. For each iteration of the permutation test, the strongest PLV of the extracted source pairs was taken as the PLV of that iteration. Finally, the NID components (extracted from the non-permuted components), whose PLVs were larger than at least 95% of the PLVs of the permutation iterations were kept as significant components.

## 4. Experimental Data

### 4.1. Simulated EEG

We used realistic head modeling to simulate EEG, consisting of cross-frequency coupled sources and additive pink noise. In these simulations, the strength of the coupling, the number of cross-frequency coupled pairs, and their mixing patterns were known a-priori; thus, allowing a calculation of the PLV of extracted source signals and errors for the extraction of activation patterns.

64-channel EEG signals were simulated based on a three-compartment realistic head model (Nolte and Dassios, 2005), with channel positions corresponding to the standard positions of EEG on the Montreal Neurological Institute head (Evans et al., 1994). The sources were modeled as multiple current dipoles located in the triangularly tessellated cortical mantle. The spatial direction and location of the dipoles were chosen randomly. For additive noise, 100 mutually independent pink noise sources were used. *N* independent coupled pairs of oscillations were generated based on the type of interaction. Unless it is mentioned otherwise, two pairs of coupled oscillations (*N* = 2) were produced. In the rest of the paper, we continue with phase-phase coupled sources; however, comparable results were achieved for amplitude-amplitude coupling.

For phase-phase coupled pairs (with PLV=1) at *f*_*n*_ = *nf*_*b*_ and *f*_*m*_ = *mf*_*b*_, a narrow-band signal centered at *f*_*b*_ was produced by band-pass filtering an array of white-Gaussian noise. The phases of the sources at frequencies *f*_*n*_ and *f*_*m*_ were obtained by frequency-warping (Nikulin et al., 2012) of the phase of the signal at *f*_*b*_, meaning that the phase of the oscillation at *f*_*b*_ was multiplied by *n* and *m*. For each of the signals at *f*_*n*_ and *f*_*m*_, the amplitude envelope was set equal to the envelope of an independent array of band-passed white-Gaussian noise at the same frequency band. We set *f*_*b*_ = 10Hz in the simulations with the motivation of having *α* band as the base frequency. The cuttoff frequencies of the band-pass filter was 8Hz and 12Hz.

Note that, a fourth-order Butterworth filter was used in all cases of band-pass filtering, applied backward-forward to prevent phase distortion. Additionally, the sampling frequency was set to 500Hz.

#### 4.1.1. Evaluating NID using simulated data

##### Evaluation criteria

For each simulation, the dissimilarity between the original, a-priori known (**p**) and the extracted 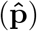 mixing patterns was measured using the following index for each extracted oscillation:

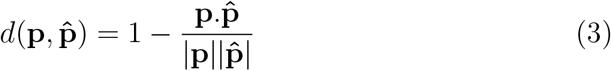

Figure 3 shows an example of a simulated activation pattern and three other patterns along with their dissimilarities to the simulated one.

**Figure 3:**
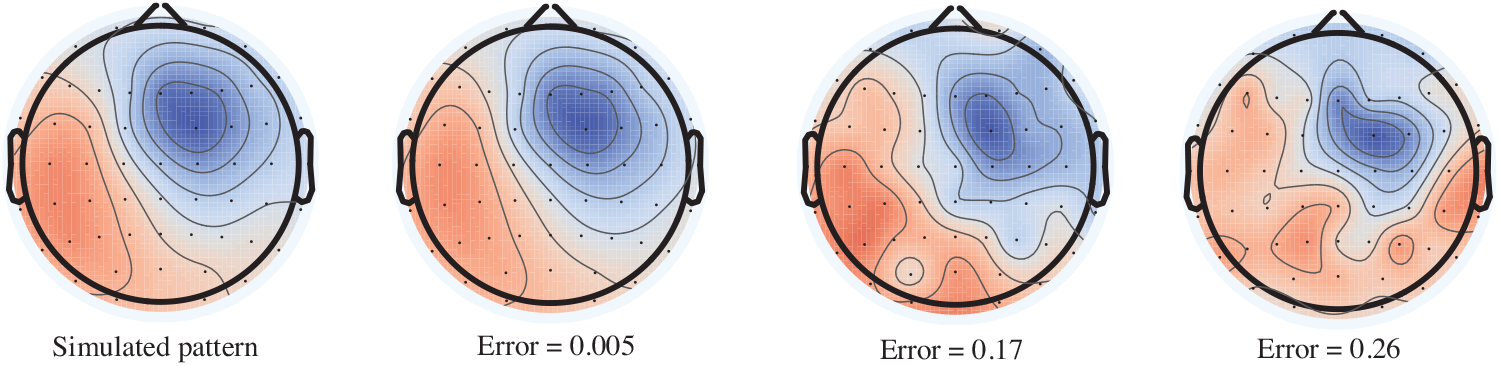
A simulated activation pattern and three other patterns and their dissimilarities (error) to the simulated one. The error is computed according to equation 3.

Since the estimated mixing patterns are compared with the ground truth when working with the simulated data, the above dissimilarity index is called the error of mixing patterns. Having *N* pairs of coupled sources in each simulation, 2*N* errors are computed. The median of these errors was reported as the representative error of the source recovery.

Another parameter that helps to evaluate the performance of the algorithm is the PLV of the extracted sources. For each simulation, the mean PLV of all extracted pairs of sources (average of *N* values) was reported.

##### Evaluation conditions

NID’s performance was examined at various signal-to-noise ratios (SNR), values for strength of coupling, and number of pairs of coupled oscillations. For each condition, 100 five-minute EEG signals were simulated comprising pairs of coupled oscillations at frequencies (10Hz,20Hz), (10Hz,40Hz), and (20Hz,30Hz), which we refer to as 1:2, 1:4, and 2:3 coupling, respectively. SNR was defined as the ratio of the mean power of the projected oscillations to the power of projected pink noise at each frequency.

Another important issue was to investigate whether the method overfits the data when finding the coupled sources. We checked this for NID considering two aspects: frequency specificity and noise overfitting. This is explained in the next two paragraphs.

Firstly, we investigated how NID performs in separating coupled source signals with the frequency ratio of *n* : *m* when the algorithm’s parameters are not set equal to n and m. To verify this, coupled sources with the frequency ratio of 1:4 were simulated, while the frequency ratio parameters of NID was set to 1:2.

Additionally, we investigated whether the algorithm overfits the noise of the data by extracting spurious sources. For this purpose, the EEG signal was first simulated without any oscillations being added (i.e. the EEG channels contained only projected noise). Second, NID was applied on the simulated EEG consisting of pink noise and two independent oscillations at each of two frequency bands of interest, i.e. [8,12]Hz and [16,24]Hz. The frequency ratio parameters of NID were then set to 1:2. The performance of NID was evaluated in 100 simulations where NID was applied to find two 1:2 coupled sources. The significance of the coupling of the extracted sources was assessed through the statistical testing described in section 3.5.

##### NID for triplets of sources

To test the reliability of NID for recovering triplets of the coupled sources (refer to section 3.4), two triplets of 1:2:3 coupled sources (oscillations at 10Hz, 20Hz, and 30Hz) were simulated at different SNRs and the performance of NID was evaluated by assessing the error of mixing patterns and the mean PLV of the extracted sources. Note that for each set of coupled oscillations, the PLV is computed as the mean PLV of 1:2 and 1:3 coupled signals.

### 4.2 Real EEG

We applied NID to resting-state EEG data from the open access database of the LEMON study (the Leipzig study for mind-body-emotion interactions). This study was carried out in accordance with the Declaration of Helsinki and the study protocol was approved by the ethics committee at the medical faculty of the University of Leipzig (Babayan et al., 2018). The dataset includes 16-min, 62-channel resting-state EEG recordings, which consist of 16, interleaved, one-minute blocks of eyes-closed (EC) and eyes-open (EO) conditions. The EEG was recorded with a band-pass filter between 0.015 Hz and 1 kHz and a sampling frequency of 2500 Hz.

For our analysis, we have used the recordings of young (20-35 years old), right-handed men, which totaled 81 subjects. From the total EEG available (16 min), only the EC condition was used, resulting in eight-minute resting EEG data for each of the subjects.

The preprocessed EEG data from the LEMON study is publicly available in the database. In the preprocessing steps, the signal has been downsampled to 250 Hz, band-pass filtered within [1,45]Hz with a fourth-order Butterworth filter (applied backward-forward), and split into EO and EC conditions. Artifact rejection has been done through visual inspection, principal component analysis (PCA), and ICA. For more details of preprocessing procedure we refer the reader to (Babayan et al., 2018).

#### 4.2.1. Extraction of interacting sources from real data

We used NID for extracting phase-phase coupling between alpha ([8,12]Hz) and beta ([16,24]Hz) frequency bands.

For each subject, five pairs of coupled oscillations were initially extracted. The significance of the extracted sources for each subject was determined with the permutation test explained in section 3.5. From the extracted pairs of coupled source signals, those that their PLVs could survive the permutation test were kept as significant and used in the further analysis.

#### 4.2.2. Evaluating NID using real data

While decomposing real data, it is not possible to examine the validity of the extracted mixing patterns. Therefore, we , firstly, examined the PLV of the extracted source signals. In the next step, in order to inspect the relationship of the spatial location of the paired oscillations, the dissimilarity of their activation patterns was calculated using equation 3. The smaller the computed index, the more similar the activation patterns. A dissimilarity of zero would indicate that the cross-frequency interactions can be due to the presence of multiple harmonics, while non-zero dissimilarities is likely to indicate the presence of genuine interactions. Spatially distinct interactions are mostly interesting for us, because they can demonstrate remote interactions in the brain. Additionally, we investigated the relationship between the PLV of the extracted coupled oscillations and the dissimilarity between their activation patterns to assess whether the spatial location of the extracted coupled oscillations has any impact on the PLV of their coupling..

Note that with synthetic data the Ground Truth is a-priori known; therefore, equation 3 gives the error of the estimated mixing patterns. However, with real biological data this equation is used to estimate the dissimilarity of the two mixing patterns.

#### 4.2.3. Localizing the activation patterns

We localized each of the extracted components in the source space using the eLORETA inverse modelling (Pascual-Marqui, 2007) and the New York head model (Haufe et al., 2015; Huang et al., 2016) with approximately 2000 voxels. The MATLAB^©^ implementation of the eLORETA algorithm is available in MEG/EEG Toolbox of Hamburg (METH). The voxels of the head model are attributed to regions of interest (ROI) based on the Harvard-Oxford atlas, which has 96 cortical ROIs. In order to analyze the relationship of the localization of alpha and beta sources at the group level, we pooled all the extracted oscillations of all subjects together. In the inverse model of each source, the voxel values were thresholded by 95% of the maximum activity across all voxels. Therefore, for the inverse model of the i-th pair, *N*_*i,α*_ and *N*_*i,β*_ voxels remained with non-zero values for the *α* and *β* oscillations respectively, from which 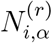 and 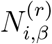 voxels were in ROI *r*.

In order to quantify the activity in each ROI, the following value was computed for all ROIs (*r* = 1, · · · , 96):

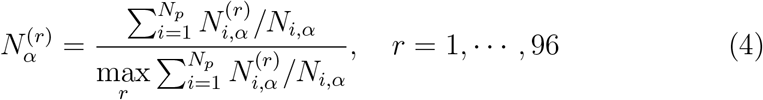

where *N*_*p*_ is the total number of extracted pairs for all the subjects. Similar equation is used to compute 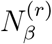 by replacing all the *α* indexes by *β*. 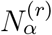, computed in equation 4, reflects the total amount of *α* activity in the r-th ROI, which is related to the number of active voxels in this ROI in all the activation patterns of all subjects.

To investigate how ROIs interact with each other, a 96×96, non-symmetric matrix **R** was calculated. Element **R**(*r*_1_, *r*_2_) of the matrix reflects the amount of interaction between *α* oscillations in ROI *r*_1_ and *β* oscillations in ROI *r*_2_. **R**(*r*_1_, *r*_2_) is not a measure of the strength of the interaction (PLV) but how often *β* activity is observed in ROI *r*_2_ when there is *α* activity in ROI *r*_1_. The following equation is used to compute **R**(*r*_1_, *r*_2_):

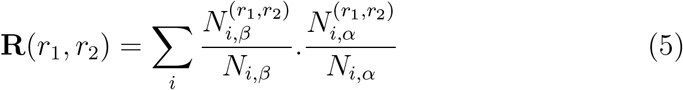

where 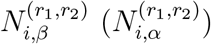 for the i-th *α*-*β* pair is the number of voxels with *β* (*α*) activity in ROI *r*_2_ (*r*_1_) when the coupled *α* (*β*) oscillation has activity in ROI *r*_1_ (*r*_2_).

**R** is an asymmetric adjacency matrix of a graph. For the visualization purposes, we converted it to a bipartite graph, which has 96 nodes in each part. It means that the (*r*_1_, *r*_2_) element of **R** translates to the edge between node *r*_1_ of part 1 and node *r*_2_ of part 2.

## 5. Results

In most cases, box-plots are used for reporting the results. The band displayed within each box is the second quartile (the median), and the box expands between the first and the third quartiles. The whiskers have a maximum length of 1.5 times the Interquartile range (IQR). Note that all the analysis were performed in MATLAB^©^ R2017b.

### 5.1. Simulations

#### 5.1.1. NID has reliable performance at different SNRs

The simulations were performed with SNR = −15, −10, −5, 0 (dB) for two coupled pairs of sources. Figure 4 depicts the error box-plots of mixing patterns and the graph of the mean PLV of the extracted sources. One can see that the median error is < 0.05 and the mean PLVs are > 0.1 even for the very low SNR of −10 (dB) (meaning that the power of noise is 10 times larger the power of the signal of interest). This shows that in the most simulation runs NID successfully recovers the activation patterns of the components. Comparable results were achieved for oscillations with amplitude-amplitude coupling: at SNR −15 (dB) with median errors of < 0.03 and mean PLVs of > 0.4 (figure S3 of supplementary material).

**Figure 4:**
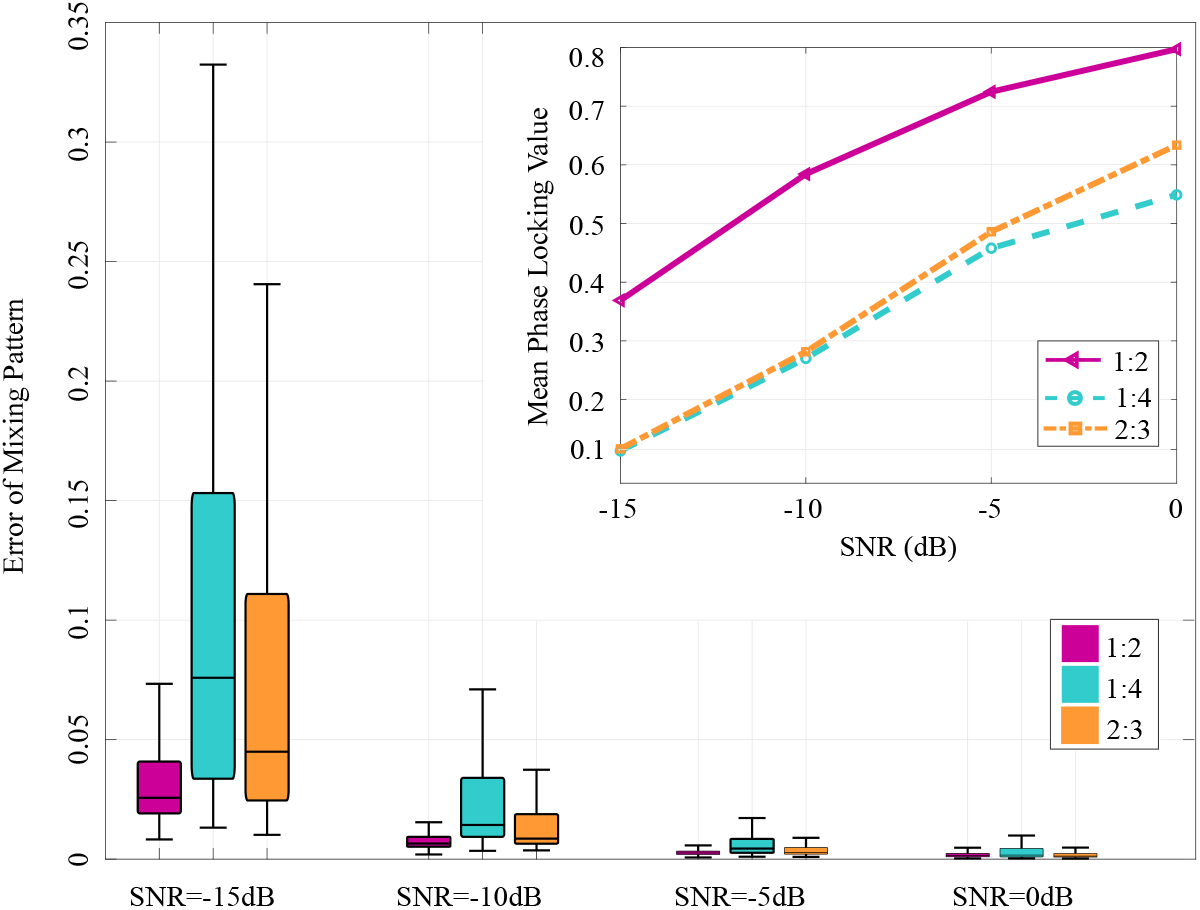
The performance of NID with simulated EEG, for the extraction of two pairs of cross-frequency phase-phase coupled oscillations at different SNRs and frequency ratios. Main plot: Box-plots of errors of mixing patterns. Subplot: Mean PLV vs. SNR for extracted components.The small median errors as well as the relatively large mean PLVs show that the performance of NID in untangling the coupled source signals is reliable.

#### 5.1.2. NID can extract multiple pairs of coupled sources reliably

We also investigated the impact of the number of interacting pairs on the performance of NID. In the previous section, two pairs of source signals were simulated. Here, five independent pairs were simulated. The box-plots of the errors of mixing patterns and mean PLVs of the extracted oscillations at SNR=−10 (dB) are illustrated in figure 5. The median errors of < 0.05 indicate that NID is successful in extracting even five pairs of interacting oscillations even at a low SNR of −10 (dB).

**Figure 5:**
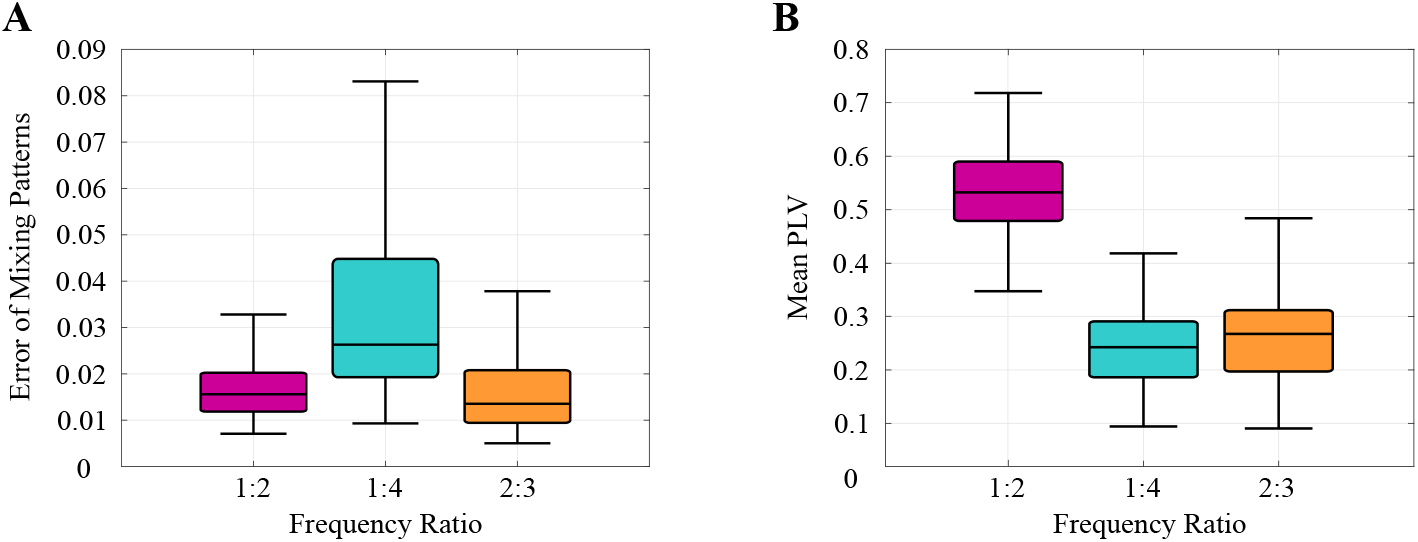
The performance of NID for extraction of five pairs of cross-frequency phase-phase coupled oscillations at SNR= −10 (dB) and different frequency ratios applied on simulated EEG. (a) Box-plots of error of mixing patterns. (b) Box-plots of the mean PLV. We conclude the reliable performance of NID from the small activation pattern errors and large mean PLVs.

#### 5.1.3. NID can extract weakly coupled oscillations

In this section, the simulations were performed for different synchronization strength at SNR=−10 (dB). Details of generating coupled sources with different synchronization strength are presented in section 2.1 of the supplementary material.

Figure 6 depicts how median errors of mixing patterns change with mean PLV of the underlying coupled sources. It is clear that even for very weak couplings, NID successfully recovers patterns of the interacting components with the corresponding errors being < 0.05.

**Figure 6:**
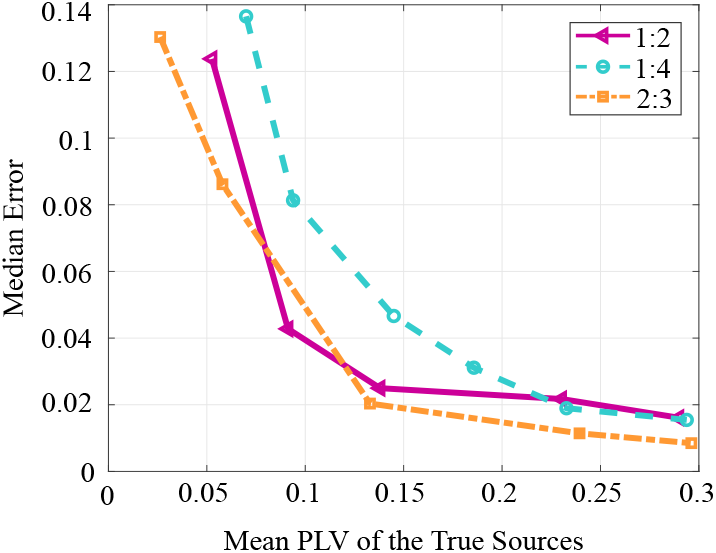
Median error vs. coupling strength (PLV) for different frequency ratios at SNR=−10 (dB) with two pairs of phase-phase coupled sources. The median errors decrease as the PLV of the simulated coupled oscilaltions increases. Even for weak couplings (e.g. PLV=0.1), the errors of pattern extraction are small.

#### 5.1.4. NID is able to detect triplets of coupled sources

As mentioned in section 3, NID can easily be generalized to extract *n* : *m*: *p* coupling (triplets of coupled sources). Figure 7 shows the box-plots of errors of mixing patterns and mean PLVs of the extracted sources when two triplets of 1:2:3 (with base frequency of 10Hz) coupled sources exist in the simulated EEG. NID can extract the oscillations reliably even at SNR=−15 (dB) with median error < 0.05 and mean PLV of the extracted sources > 0.3.

**Figure 7:**
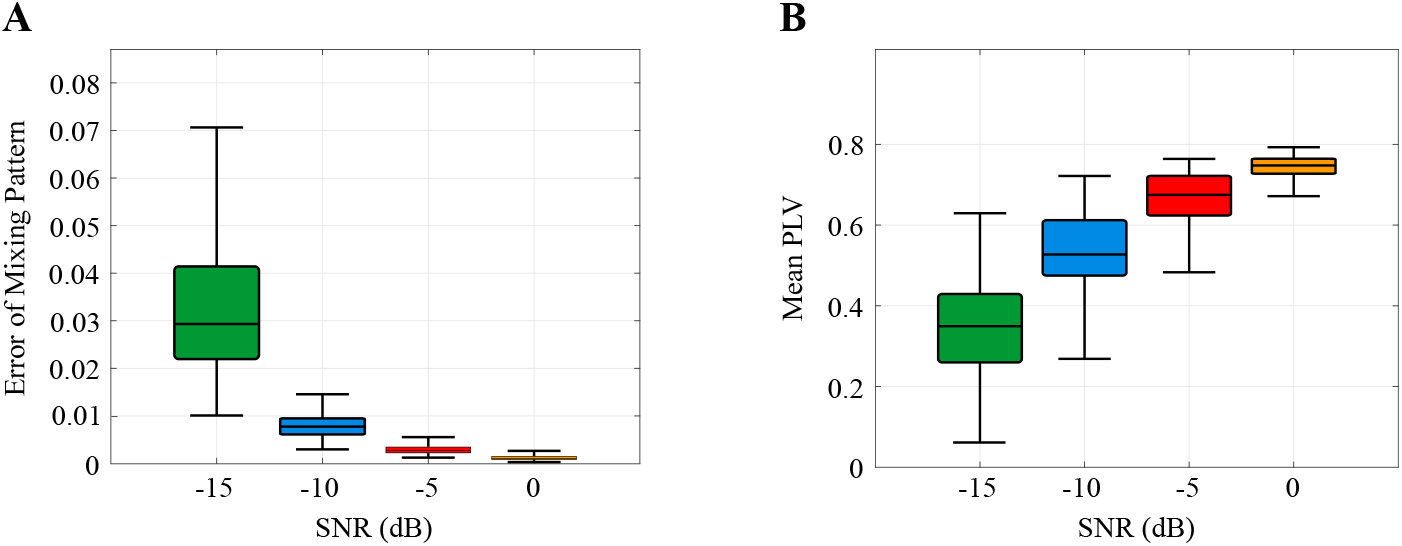
The performance of NID for the extraction of two 1:2:3 coupled triplets of oscillations at different SNRs. (A) Mixing pattern error box-plots. (B) Mean PLV of the extracted sources. NID extracted the source signals reliably.

#### 5.1.5. NID does not overfit the data

As mentioned in section 4.1.1, we investigated the behavior of NID in the case where the frequency ratios are specified with mismatches, or where there are no coupled sources in the data.

For checking the frequency specificity, 1:4 phase-phase coupled sources were simulated at SNR=−10 (dB), while NID’s frequency ratio parameter was set to 1:2. Such mismatch in frequencies lead to very large errors (median error> 0.3, supplementary figure S2), indicating that the successful extraction of the coupled components requires a-priori knowledge of frequency information.

In addition, NID was applied to simulated EEG consisting of only noise, or with 2 uncoupled sources at each of the frequencies of interest. In the former case, the median of the PLV of the extracted pairs was 0.025, and non of the pairs of the extracted sources survived permutation test (Bonferroni multiple testing corrected).

### 5.2. Resting-state EEG

With the procedure explained in section 4.2.1, a total number of 243 alpha-beta, significant, interacting pairs of oscillations were extracted from all the subjects.

### 5.3. NID-component analysis

As mentioned in section 4.2.2, the dissimilarity between the mixing patterns of each pair of source signals was computed using equation 3. Figure 8-A illustrates the relation between the strength of coupling and the similarity of the activation patterns of each pair. No significant linear correlations was observed between these two variables. Additionally, figure 8-B shows a box-plot of the PLVs of the extracted pairs of oscialltions. Comparing these PLVs with the PLVs in the sensor-space, one can clearly see a two-fold improvement in the estimation of PLV using NID. A box-plot representation of the sensor-space PLVs (median of 0.06) of the subjects is presented in supplementary figure S6.

**Figure 8:**
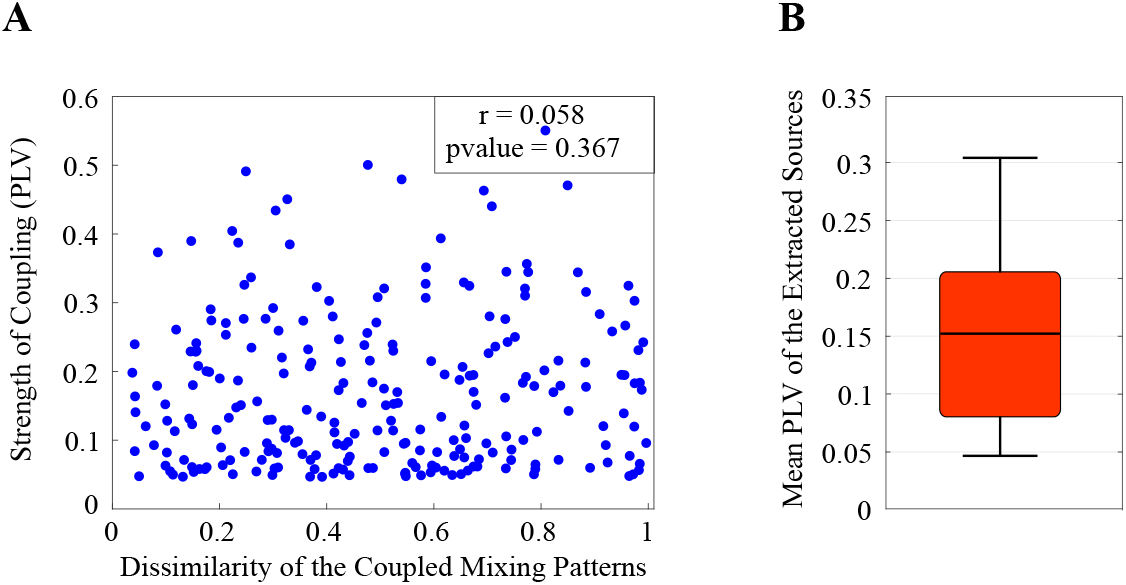
(A) The relation between PLVs of each pair and dissimilarities between their mixing patterns. (B) Box-plots of the PLVs of extracted phase-phase coupled oscillations.

### 5.4. Localization of NID components

Referring to section 4.2.3, figure 9 illustrates the ROI-based localization of NID components using the values calculated with equation (4). For both frequencies, subjects have non-linearly interacting sources primarily in occipital regions extending to parietal regions, as well as in the sensorimotor areas extending to the frontal regions. Additionally, beta activity in the sensorimotor areas occurred more frequent than alpha activity.

**Figure 9:**
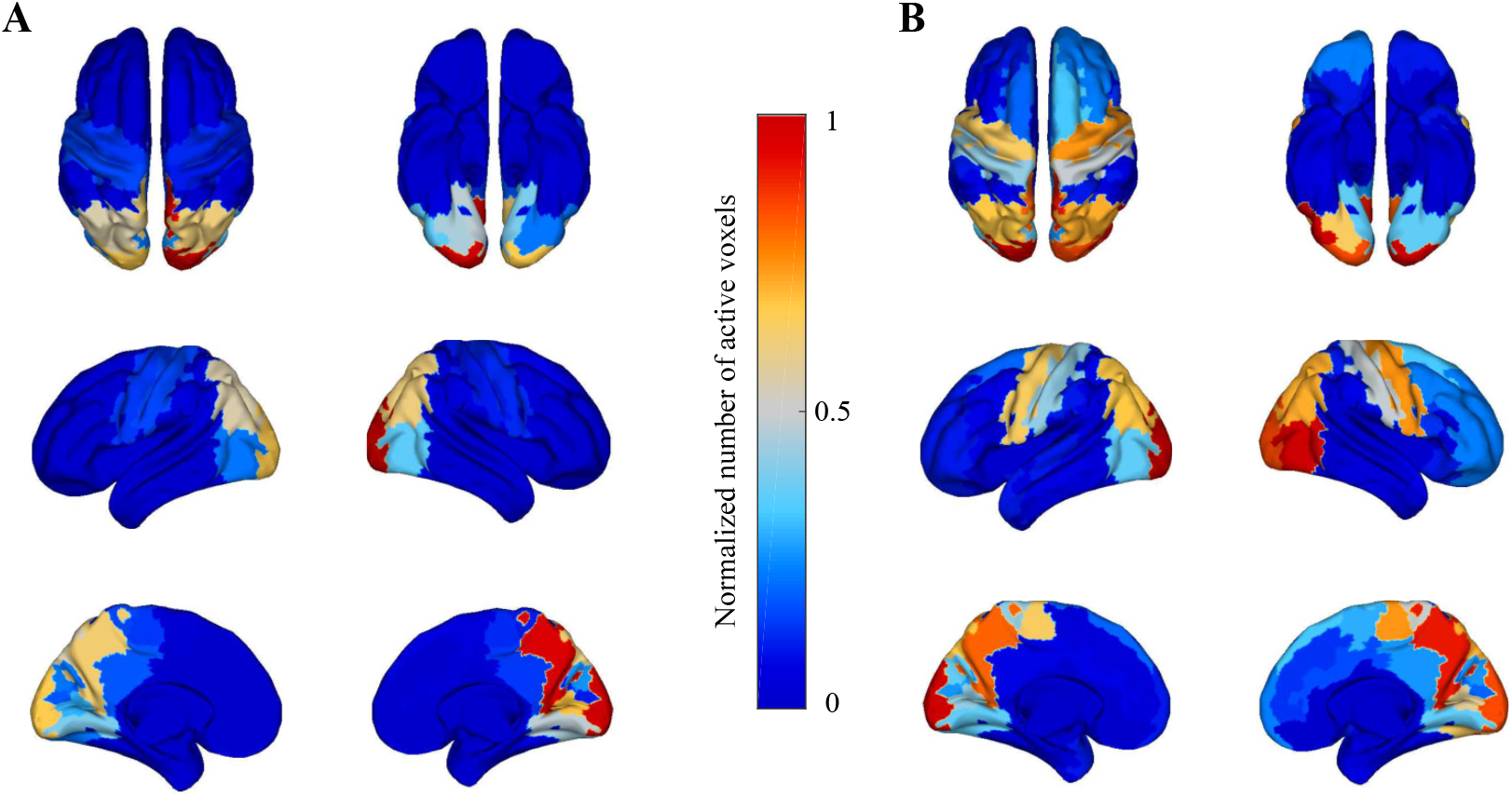
ROI-based analysis of NID source signals based on equation 4 for (A) alpha and (B) beta oscillations. For both frequencies, subjects have mainly activities in occipital, parietal, sensorimotor, and frontal regions. The beta-activity in motor regions are more pronounced than alpha-activity.

Computed in equation 5, we have a measure of the interactions between different brain regions, which is depicted in figure 10. These interactions can be depicted with a weighted, bipartite graph, whose nodes are the ROIs and where the edges denote the interactions between two ROIs. The connection between nodes *r*_1_ of the upper part and *r*_2_ of the other part indicates that there is alpha-activity in ROI *r*_1_ that is interacting with beta-activity in ROI *r*_2_. The weight of the edges are proportional to the number of active voxels in the two regions. Figure 10-A depicts the bipartite graph representing the adjacency matrix of ROI-interactions. (pre-)Frontal areas, and pre- and post-central gyri of both hemispheres have beta sources which interact with alpha sources of other ROIs. Additionally, in precuneous cortex and occipital areas both alpha- and beta-sources have interactions with the sources of alpha and beta oscillations at multiple ROIs. Some medial ROIs show interactions for their beta-sources in one of the hemispheres. The most connected regions in figure 10 can also be observed in figure 9.

**Figure 10:**
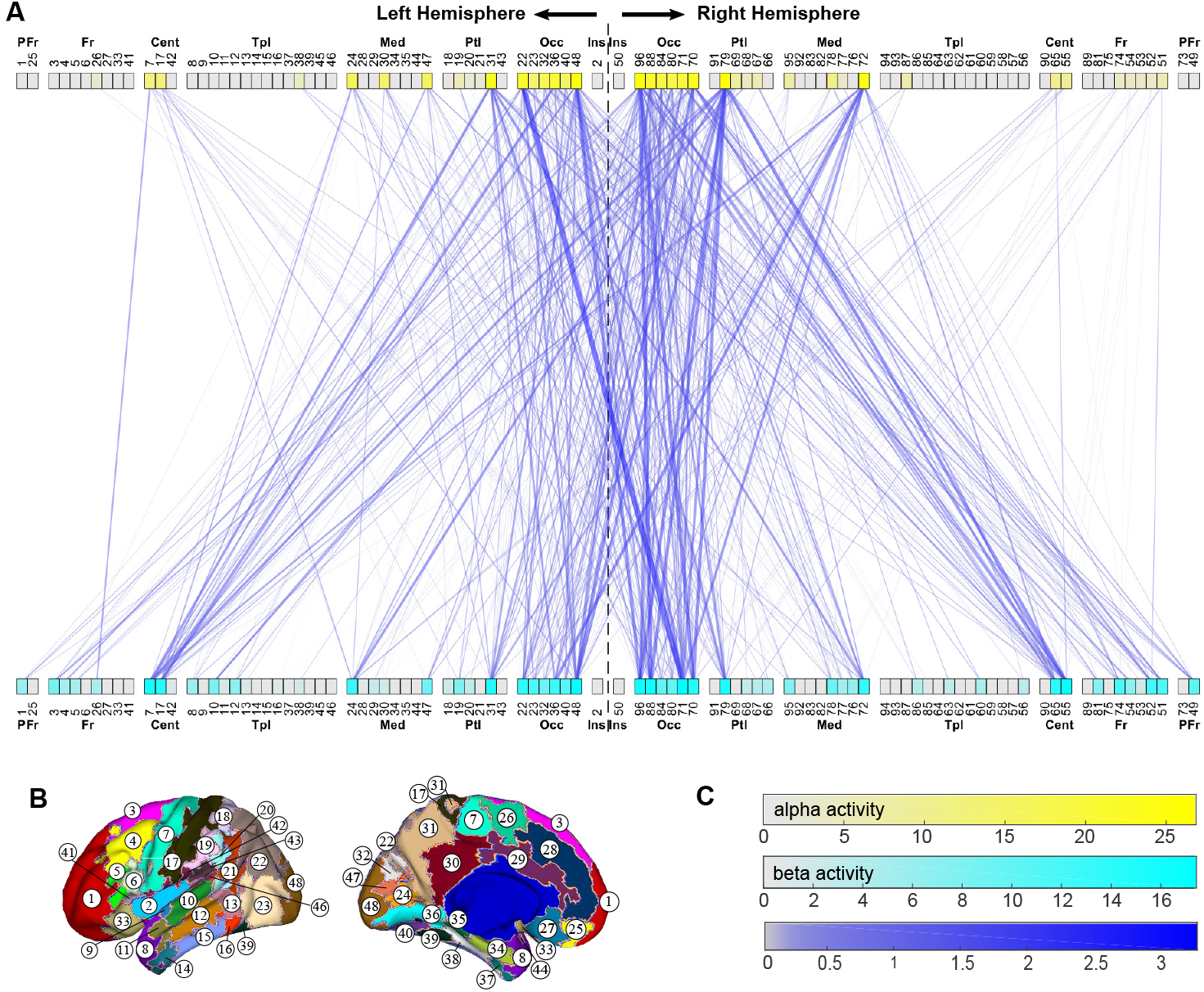
The bipartite graph illustrating the alpha-beta interactions among ROIs of Harvard-Oxford atlas, computed for 82 subjects. A connection between node *r*_1_ of the upper and node *r*_2_ of the lower part indicates that alpha oscillations in ROI *r*_1_ interact with beta-oscillations in ROI *r*_2_; the weight of the connecting edge is proportional to the number of active voxels (across subjects) in the two ROIs. The weight of an edge is coded by the width and color-strength of the edges. The degree of the nodes are color-coded. Panel B depicts the Harvard-Oxford atlas ROIs of left hemisphere, and panel C depicts the color-codes.

## 6. Discussion

We introduced a novel, general framework for the extraction of cross-frequency coupled sources from EEG/MEG, namely Non-linear Interaction Decomposition (NID). The idea of assessing the distribution of a mixture of coupled oscillations is introduced for the first time and provides a novel perspective for investigating non-linear interactions in EEG/MEG.

We validated the method with extensive simulations in different conditions. NID showed reliable performance in the extraction of cross-frequency phase-phase and amplitude-amplitude coupled oscillations in simulated EEG even at a very low SNR of −15 (dB) and also for weak coupling strengths. We also used NID for extracting phase-phase coupled sources in human resting-state EEG data. The PLV was found to be considerably higher compared to the PLV obtained for each of the single channels, while multiple testing and uncertainty caused by volume conduction were avoided by projecting the data to the lower dimensional space of NID.

NID can be used for the extraction of coupled sources originating from different recording modalities or investigating the interactions between different subjects. NID is also generalizable to the investigation of interactions between more than two frequency bands, e.g. alpha-beta-gamma, which is not possible through other methods. In addition, the algorithm has the potential to be tuned for a specific type of coupling through the contrast function of the non-Gaussianity maximization step, although this latter aspect requires a more systematic investigation.

### Remote interactions are captured by NID

An important feature of NID is that it separates coupled oscillations at distinct spatial locations. We tested this by computing the dissimilarity between the topographies of the extracted sources where larger values indicate spatially distinct sources. It could be the case that sources which have similar topographies are harmonics of a non-sinusoidal source in the brain and therefore, we can investigate the remote interactions by assessing the dissimilarity of mixing patterns of coupled sources. The relation between the PLV of the source pairs and the dissimilarity of their topographies is plotted in figure 8, which shows that they are not linearly correlated. Therefore, one can conclude that strong interactions (high PLV) exist for sources with similar topographies as well as for those with different topographies, showing that NID is able to extract spatially distinct oscillations with large PLVs. This finding can also be observed on the bipartite graph of figure 10, which illustrates the existence of remote interactions between different ROIs. From this graph, diverse interactions between the two hemispheres, or between central, parietal, and occipital areas can be observed.

There is a rich literature focused on alpha and beta oscillations in the brain. The oscillations in the alpha-frequency range, are particularly prevalent in parietal and occipital regions, while beta-oscillations are pronounced over sensorimotor cortex (Groppe et al., 2013; Tewarie et al., 2016). In line with these observations, figure 9 shows the presence of alpha activity in occipital and beta activity in sensorimotor regions. Moreover, figure 10 suggests the existence of interactions between beta oscillations in central and alpha oscillations in occipital areas. These may be viewed as a functional substrate for visuo-motor integration (Tewarie et al., 2016). There is actually an anatomical evidence that these two areas are indirectly connected which might be important for sensory guidance of movement (Glickstein, 2000; Kravitz et al., 2011; Strigaro et al., 2015). Our results suggest that such indirect anatomical connectivity can be manifested electrophysiologically through alpha-beta phase-phase coupling in resting-state.

### 6.1. Relation to previous methods

ICA is frequently used for the extraction of EEG/MEG sources signals. Since NID has a non-Gaussianity maximization decomposition (NGMD) step, it is necessary to emphasize the distinction between NID and ICA. The main technical difference between NID and ICA is that an augmented matrix of different frequency contents is decomposed to maximally non-Gaussian components. This is in contrast to ICA methods (e.g. JADE, fastICA, InfoMax), where the broad-band multi-channel signal is decomposed. This very difference gives the NGMD algorithm the flexibility to select different weights for the components at different frequencies. Therefore, the weights of the linear mixture of the coupled sources are selected flexibly to make the mixture maximally non-Gaussian, while ICA forces all frequency contents to be mixed with the same weights. Moreover, the SNR has been improved in the two frequency bands of interest via the application of SSD in the first step of NID, which clearly contributes to its successful performance.

There are not many multivariate methods for the extraction of cross-frequency coupled sources. The novelty of NID lies in the extraction of the coupled sources based on the statistical properties of the coupled oscillations. Other methods are optimized for the detection of a specific coupling; however, they can also be sensitive to other types of coupling. As an example, our simulations show that cSPoC (Dähne et al., 2014), optimized for detection of oscillations with power dependencies, is also able to detect phase-phase coupled sources ^2^. Although it is not surprising that NID outperforms cSPoC in the extraction of phase synchronized sources, we emphasize that there is no explicit optimization of any contrast function based on the type of coupling in NID’s algorithm. NID is at least as good as cSPoC (Dähne et al., 2014) in detection of cross-frequency amplitude-amplitude coupling (figure S4 of supplementary material), while being 1.5 times faster. It is worth mentioning that it has been shown that cSPoC outperforms other methods in the extraction of oscillations with power dependencies (Dähne et al., 2014).

Cross-frequency decomposition (CFD) (Nikulin et al., 2012) is a multi-variate method for the detection of phase synchrony in MEG/EEG. While NID imposes no restriction on frequency ratios (*n* : *m* coupling, *n, m* ∈ ℕ), CFD only works for the case where n=1. The supplementary figure S5 compares the performance of the two methods in the extraction of two pairs of 1:m (*m* ∈ ℕ) phase-phase coupled sources. Although both methods have reliable performance, NID still outperforms CFD. Generalized cross-frequency decomposition (GCFD) (Volk et al., 2018) is a generalization of CFD for arbitrary frequency ratios *n* : *m* (*n, m* ∈ ℕ). GCFD extracts the phase-phase coupled sources by finding the spatial filter that optimizes the correlation of frequency-warped SSD components. This approach results in a reliable extraction of the coupled sources; however, it is asymmetric (i.e. depends on which band is used as a regressor) and computationally expensive. Additionally, frequency-warping (multiplying the phase of a signal by a factor) distorts the frequency content of a signal; therefore, the relations of frequency-warped signals may not directly reflect true oscillations in the brain.

### 6.2. Future work

Our observation is that the distribution of the mixture of cross-frequency coupled sources differs depending on the type of coupling. For example, we observed that the distribution of the mixture has longer tails for amplitude-amplitude coupled source signals, while it is skewed and has “shoulders” for phase-phase coupled sources. These properties can be better explained by different measures. For instance, “tailedness” is expressed best by kurtosis, while skewness can be described the distributions skewed to one side. Additionally, we know that higher order odd moments of a Gaussian signal are zero; therefore, they can explain some features of non-Gaussian signals. Consequently, one of the future works for extending the NID algorithm is how to define the NGMD contrast function to get even better results for different types of coupling.

In recent years, there has been a considerable interest to whole-brain connectivity and its relation to cognitive performance (Palva et al., 2010; Palva and Palva, 2012; Siebenhühner et al., 2016; Siebenhuehner et al., 2019). In this regard, brain networks demonstrating cross-frequency interactions are becoming popular as well (Siebenhühner et al., 2016; Siebenhuehner et al., 2019; Tewarie et al., 2016), reflecting the importance of spectrally distributed information processing in the brain. Using multivariate methods like NID for extraction of a subspace of brain oscillations with cross-frequency coupling, can be helpful for alleviating signal mixing problem and extracting meaningful interacting components. These components can then be used for further MEG/EEG analysis e.g. to investigate the properties of cross-frequency brain networks in resting-state or during the cognitive, sensory and motor task performance.

## Appendix A Further discussion about NID

In this section we discuss that the non-Gaussianity maximization step of the NID algorithm is able to separate the non-linearly coupled sources.

SSD components and patterns can be modeled as a mixture of the true sources and their mixing patterns according to:

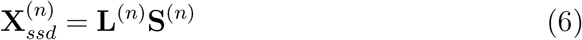

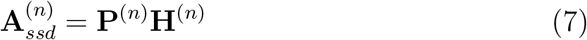

The i-th row of **H**^(*n*)^ ∈ ℝ^*N*×*N*^ includes the contribution weights of 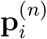 in SSD patterns. Also, the i-th column of **L**^(*n*)^ ∈ ℝ^*N*×*N*^ represents the contribution weights of 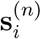 in SSD components.

Without the loss of generality, for the sake of simplicity, we assume the case of *n* : *m* coupling, i.e. coupling of pairs of sources.

Let 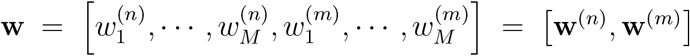 be the filter relating to one of the dimensions of NGMD subspace, then **r** = **wX**_*Aug*_ would be the projection of the SSD sources on this dimension. We can rewrite **r** as follows:

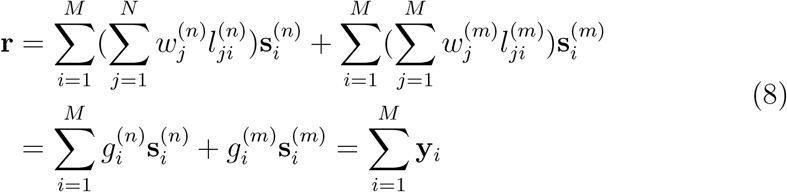

where 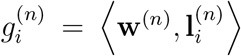, and 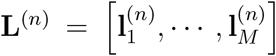. Additionally, 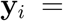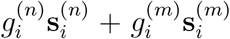 is defined as the weighted sum of signals of pair *i*. By assuming that each coupled pair is independent from other pairs, **y**_*i*_ is independent of **y**_*j*_ for *i* ≠ *j*. Since, by assumption, the sources at each frequency have approximately the same distribution, we expect **y**_*i*_, ∀*i* to be roughly identically distributed. From central limit theorem we know that the sum of i.i.d. random variables is “more Gaussian” than each of them separately. Thus, we can claim that the non-Gaussianity of **r** is maximized if it is equal to one **y**_*i*_, meaning that 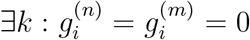, *i* ≠ *k*. This means that the *i*^*th*^ source of NGMD is the mixture of signals of the *k*^*th*^ pair.

Algorithms of ICA can be used as the non-Gaussianity maximization decomposition. In addition, any contrast function maximizing the non-Gaussianity can be exploited. For instance, we suggest the following contrast function:

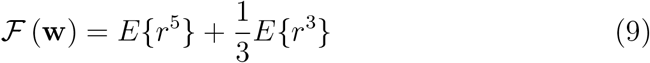

where *r* is the random variable representing **r**. 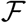 is a combination of fifth order moment of the projected signal and its skewness. It is known that the fifth order moment and skewness of a Gaussian variable are zero; therefore, by maximizing the contrast function in equation 9 we are maximizing the non-gaussianity.

Our strategy to maximize the non-gaussinity of projected sources is to take the advantage of both the contrast function in equation 9 and JADE (Cardoso and Souloumiac, 1996) algorithm. Therefore, both contrast functions (JADE and equation 9) are optimized and the optimization procedure which produces projections with maximum negentropy (maximum non-gaussianity (Hyvärinen and Oja, 2000)) is selected.

## Appendix B Practical details of computing the final mixing patterns

Each of 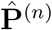 and 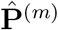 in equation 2 contain 2N patterns and N of them should be selected (i.e. N pairs of interacting sources should be selected). For this purpose, we firstly find the similar pairs; i.e. those pairs *i* and *j*, for which 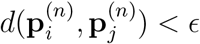 and 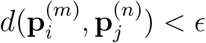, where *d*(., .) is computed as the dissimilarity between the two patterns as in equation 3. Among the similar pairs the one with largest negentropy (largest non-gaussianity (Hyvärinen and Oja, 2000)) is selected and others are omitted. Afterwards, from the remaining pairs, pairs with the largest PLV (or envelope correlation) are selected as the final mixing patterns.

## Footnotes

1 The range of frequencies in each frequency band slightly differs in different references.

2 This phenomenon that methods designed for detecting a specific coupling detect other types of couplings is also reported in the literature (Hyafil, 2015).

