## Supplementary text for "Nonlinear Interaction Decomposition (NID): A Method for Separation of Cross-frequency Coupled Sources in Human Brain"

**Abstract**

In this supplementary text, some analysis details are presented in the first and second section. Additionally, supplementary figures can be find in section 3.

### 1. Linear mixture of two coupled oscillations

In this section we show that phase-phase and amplitude-amplitude coupled narrow-band oscillations can be linearly mixed in the way that the mixture has non-Gaussian distribution. For this purpose, we first show that two oscillations $X$ and $Y$ with phase-phase coupling can be linearly mixed with

weights $k_1$ and $k_2$ such that the linear combination has non-zero skewness, kurtosis, and 5th order moment.

We consider two oscillatory random variables $X = A\cos(\Phi_x)$ and $Y = B\cos(\Phi_y)$ which represent amplitude-modulated narrow-band oscillations. $X$ and $Y$ are called 1:n phase-coupled if there is an integer $n > 1$ for which $\Phi_y = n\Phi_x = n\Phi$. Therefore, we can rewrite the two oscillations as $X = A\cos(\Phi)$ and $Y = B\cos(n\Phi)$, where $A$ and $B$ are the random variables of the amplitude-envelopes and $\Phi$ is the random variable of the phase of the slower oscillation. Since the amplitude of brain oscillations changes very slowly, one can assume that the distributions of $X$ and $Y$ are roughly Gaussian or that the distribution does not deviate strongly from a Gaussian distribution in a large sense [1]. As a result of this assumption, the third and fifth order moments of $X$ and $Y$ can be estimated as zero. Without the loss of generality, we assume that $X$ and $Y$ have unit variance. Thus, their fourth moment can be approximately estimated as three (although it tends to be larger than three [1], this does not affect our final conclusion.). Using the power-reduction formulas, one can calculate the r-th moments ($r = 3, 5$) and kurtosis of a linear mixture of $X$ and $Y$ like $k_1X + k_2Y, k_1, k_2 \in \mathbb{R}$ as the following (reminding that the expected value of amplitude modulated cosine is zero):

$$E\{(k_1X + k_2Y)^3\} \approx 3k_1^2k_2E\{X^2Y\} = \frac{3}{4}k_1^2k_2E\{a^2b.cos((n-2)\Phi)\}$$
$$= \begin{cases} \frac{3}{4}k_1^2k_2E\{a^2b\}, & \text{if } n = 2 \\ 0, & \text{otherwise} \end{cases} \tag{1}$$

$$\begin{aligned} E\{(k_1X + k_2Y)^5\} &\approx 5k_1^4k_2E\{X^4Y\} + 10k_2^2k_2^3E\{X^2Y^3\} \\ &= \frac{5}{4}k_1^4k_2E\{a^4b.cos((n-2)\Phi)\} + \frac{5}{16}k_1^4k_2E\{a^4b.cos((n-4)\Phi)\} \\ &+ \frac{15}{2}k_1^2k_2^3E\{a^2b^3.cos((n-2)\Phi)\} \\ &= \begin{cases} \frac{5}{4}k_1^4k_2E\{a^4b\} + \frac{15}{2}k_1^2k_2^3E\{a^2b^3\}, & \text{if } n = 2 \\ \frac{5}{16}k_1^4k_2E\{a^4b\}, & \text{if } n = 4 \\ 0, & \text{otherwise} \end{cases} \end{aligned} \tag{2}$$

$$
\begin{aligned}
kurtosis(k_1X + k_2Y) &= E\{(k_1X + k_2Y)^4\} - 3E^2\{(k_1X + k_2Y)^2\} \\
&= k_1^4E\{X^4\} + k_2^4E\{Y^4\} + 4k_1^3k_2E\{X^3Y\} + 6k_1^2k_2^2E\{X^2Y^2\} \\
&- 3\left(k_1^2E\{X^2\} + k_2^2E\{Y^2\}\right)^2 \\
&\approx \begin{cases} 3k_1^2k_2^2\left(\frac{1}{2}E\{a^2b^2\} - 2\right) + \frac{1}{2}k_1^3k_2E\{a^3b\}, & \text{if } n = 3 \\ 3k_1^2k_2^2\left(\frac{1}{2}E\{a^2b^2\} - 2\right), & \text{otherwise} \end{cases}
\end{aligned} \tag{3}
$$

From the above equations, one can conclude that the third and fifth order moments are nonzero for the case of $n = 2, 4$ and the multipliers $k_1$ and $k_2$ can be tuned so that kurtosis is nonzero for $n = 3$. For the case of $n \neq 3$, the kurtosis is zero only if $E\{a^2b^2\} = 4$.

Similar equations can be rewritten for other cases of $n : m$ coupling (i.e. $m\Phi_x = n\Phi_y$) and it can be shown that for all cases $kurtosis(k_1X + k_2Y) = 3k_1^2k_2^2\left(\frac{1}{2}E\{a^2b^2\} - 2\right)$, which is nonzero if $E\{a^2b^2\} \neq 4$. Additionally, for the cases when $n + m$ is odd, the (n+m)-th order moment is nonzero, which is a measure of non-Gaussianity.

The above equations also hold for the case where $X$ and $Y$ are amplitude-amplitude coupled.

### 2. Analysis Details

#### *2.1. Simulation of sources with weak interaction*

To control the strength of coupling in section 5.1.3 of the main manuscript, a random variable with von Mises distribution is added to the phase of the signal at $f_m$ to provide a jitter for phase alignmnet betwen the two processes. The parameter $\kappa$ of the von Mises distribution is the parameter to control the level of phase locking. For $\kappa = 0$ von Mises distribution is equivalent to uniform distribution, and for larger $\kappa$ it has a bell-shaped distribution with $1/\kappa$ to be a measure of the concentration.

Figure S1 depicts the von Mises distribution of phase-jitter for some values of $\kappa$. As it can be seen, increasing $\kappa$ causes the phase-jitter to be restricted to a small range, resulting in a stronger phase locking. For $\kappa = 0$ there is no phase locking between the two signals.

### 3. Supplementary Figures

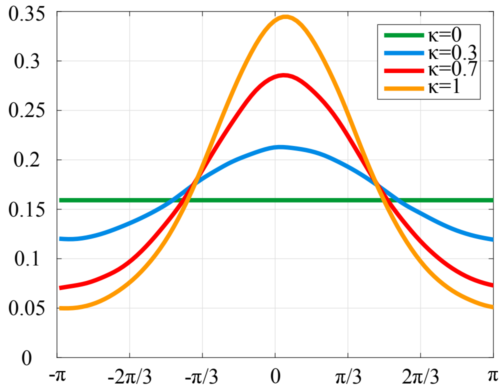


Figure S1: The pdf of von Mises distribution with mean zero and different $\kappa$. Larger $\kappa$ results in a smaller variance of the distribution.

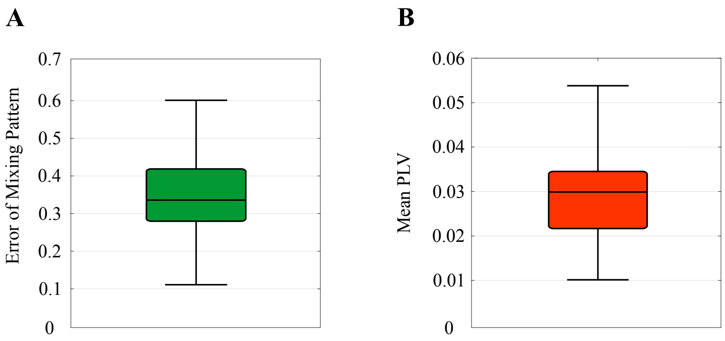


Figure S2: Error box-plot of the case when 1:4 oscillations at SNR=$-10$ (dB) are simulated and NID with parameters 1:2 is used to separate the coupled source signals. The large errors of pattern extraction (A) and small PLVs (B) show that NID fails to extract the sources if the frequency ratio parameters are not set correctly.

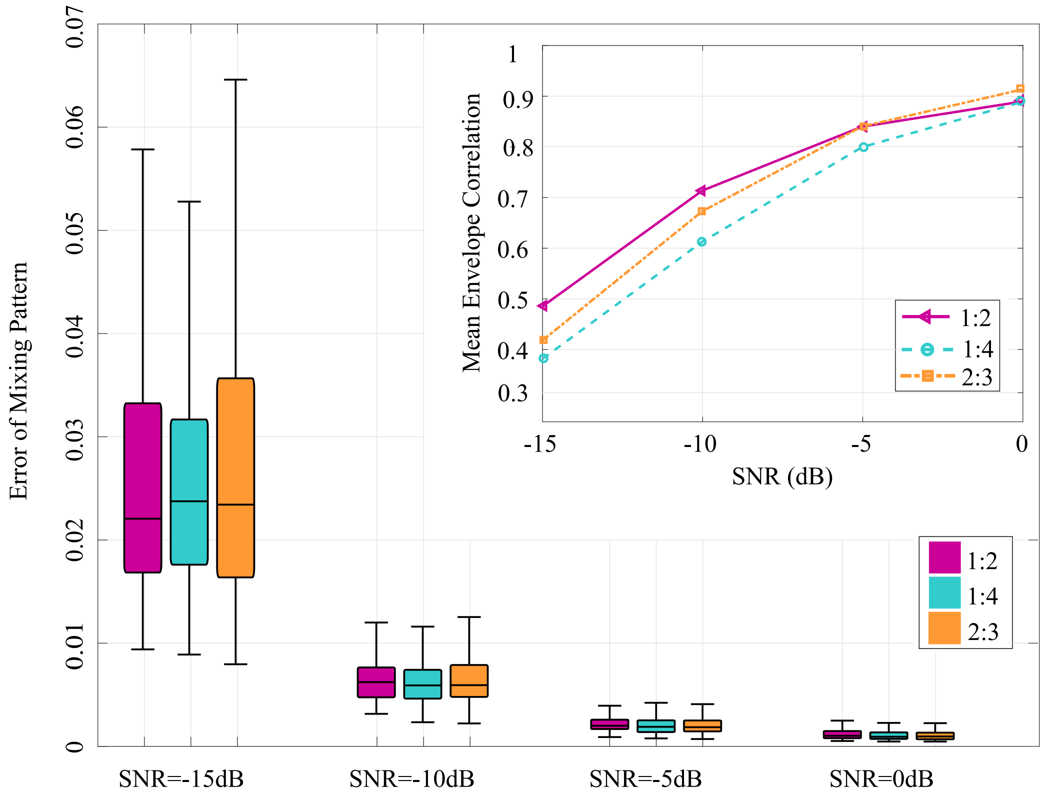


Figure S3: Performance of NID for the extraction of two pairs of cross-frequency amplitude-amplitude coupled oscilaltions at different SNRs and frequency ratios. Main plot: Errors box-plots of mixing pattern extraction. Subplot: Mean envelope correlation vs. SNR for extracted components. From the small median errors as well as the large mean envelope correlations show that the performance of NID in untangling the coupled sources is reliable.

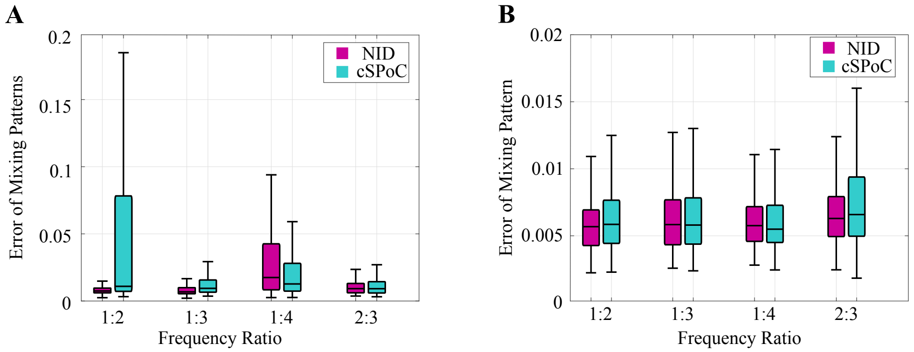


Figure S4: Comparison of the performance of cSPoC and NID in extraction of cross-frequency (A) phase-phase and (B) amplitude-amplitude coupling at SNR=−10 (dB) with two pairs of coupled oscillations. NID is at least as good as cSPoC in all cases, while it performs better in the some cases of phase-phase coupling.

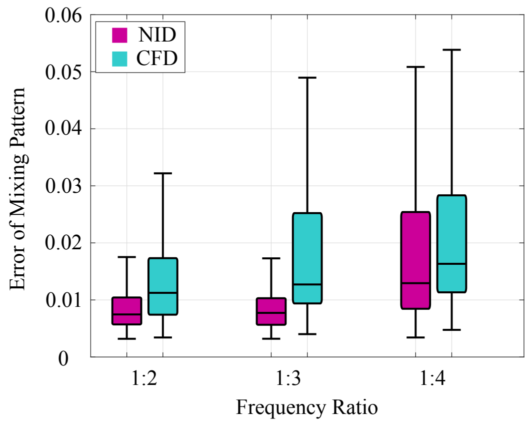


Figure S5: Comparison of the performance of CFD and NID in extraction of 1:n ($n \in \mathbb{R}$) cross-frequency phase-phase coupling at SNR=−10 (dB) with two pairs of coupled oscillations. NID outperforms CFD.

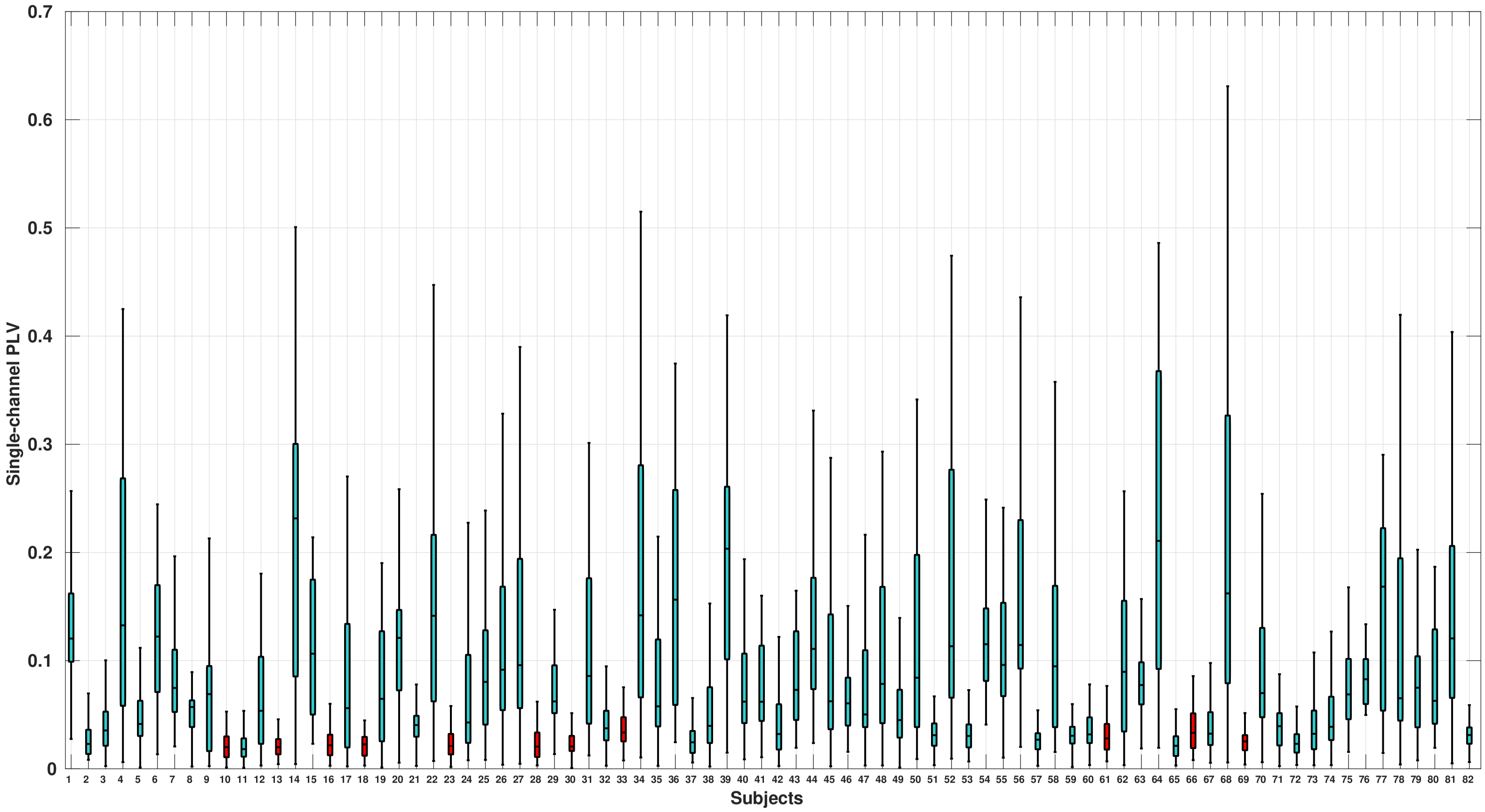


Figure S6: Box-plots of the sensor-space single-channel PLVs for all 82 subjects of LEMON dataset. Red is used for those subjects whose NID components could not pass the permutation test. One can see that for these subjects no large PLV is observed in any of the single-channels.
